# ProtJEPA: A Multimodal Joint-Embedding Predictive Architecture for Protein Biological World Modeling with Multi-Teacher Modality-Attentive Fusion

**DOI:** 10.64898/2026.08.03.742606

**Authors:** Vaibhava Lakshmi Ravideshik, Jinha Kim, Manolis Kellis

**Affiliations:** Massachusetts Institute of Technology, Computer Science and Artificial Intelligence Laboratory (CSAIL)

**Keywords:** protein embeddings, joint-embedding predictive architecture, JEPA, world modeling, multimodal learning, protein language models, knowledge graphs, self-supervised learning, biological representation learning

## Abstract

Over 99.9% of known protein sequences lack experimentally validated functional annotations. We present ProtJEPA, a multimodal Joint-Embedding Predictive Architecture that trains a sequence-only student encoder to predict joint embeddings spanning ten biological modalities—sequence, structure, knowledge graph, protein interactions, literature, localization, tissue expression, GO function, anatomy, and disorder—requiring only sequence at inference. The key innovation is target whitening, which eliminates severe anisotropy in joint targets (mean cosine 0.984 to 0.086) and prevents representation collapse without covariance regularization. On 1,828 held-out dark proteins with zero primary Pfam family overlap with training, ProtJEPA achieves 58.07% Hit@10 on zero-shot GO retrieval (+2.80 pp, ***p* = 0.020**), 69.99% enzyme class accuracy (+9.64 pp, ***p <* 0.001**), and +11.87 pp subcellular localization at 1% labels (***p <* 0.001**). Under realistic dark-protein deployment conditions where relational modalities are unavailable, ProtJEPA significantly outperforms naive concatenation of remaining modalities. Cross-domain evaluations on drug–target interaction and disorder prediction confirm transfer beyond training modalities, with the T1-only ***<*** ESMC ***<*** ProtJEPA ordering replicated across six independent tasks. Ablations establish that Phase 1 aggregator pretraining and target whitening are each independently load-bearing.

## 1 Introduction

### The protein annotation crisis

The UniProt KnowledgeBase contains over 250 million protein sequences, yet fewer than 0.1% carry experimentally validated annotations. Every unannotated protein is a potential drug target, disease mechanism, or biological pathway component that remains invisible to researchers. Current computational approaches fail to close this gap because they are fundamentally unimodal: protein language models such as ESM-2 and ESM-3 encode evolutionary information but operate solely on sequence; graph neural networks on knowledge graphs such as PrimeKG capture relational context but ignore intrinsic biochemistry; text-based approaches using BioBERT extract curated functional knowledge but cannot generalize beyond what has been written. None integrates what a trained biologist does: combining sequence conservation, structural topology, interaction context, disease associations, and textual knowledge simultaneously.

### The modality trap

We identify a fundamental limitation we term the **modality trap**: current methods over-allocate representational capacity to their training modality while remaining blind to cross-modal functional logic. A protein’s function emerges from the intersection of sequence, structure, and context simultaneously—no single-modality model can encode this correctly.

### JEPA as biological world modeling

LeCun’s World Model framework proposes that intelligence emerges from predicting abstract consequences in latent space, not from reconstructing observations. ProtJEPA operationalizes this for protein biology: given a sequence, predict complete biological context. ProtJEPA advances protein representation learning on several fronts: it is the first multimodal biological JEPA framework integrating ten complementary teachers; it introduces target whitening as a batch-size-independent anti-collapse mechanism; it establishes the Dark Protein Zero-Shot Experiment as a falsifiable benchmark for cross-modal transfer; and it demonstrates that distilled representations outperform concatenation of all available teacher embeddings under realistic dark-protein deployment conditions.

## 2 Methods

### Architecture

ProtJEPA adopts the SALT-JEPA framework [1] with ten independently pretrained, frozen teacher encoders (Supplementary Section B) fused by the MT-MA-JEPA aggregator via sample-dependent gating: *α_m_* = softmax(*q^⊤^* tanh(*W_m_h_m_*)), *z*_joint_ = LayerNorm(Σ*_m_ α_m_h_m_*) (Fig. 1). The student encoder (ESMC 600M, 585.9M trainable parameters) receives only masked protein sequence and predicts the joint embedding via a 3-layer predictor head (*d*_out_ = 512). Ten frozen teachers span sequence (ESMC 6B), structure (ESMFold2), knowledge graph (HGT on PrimeKG), PPI (GATv2), literature (BioMedBERT-large), localization, tissue expression, GO function, anatomy, and disorder; full specifications are in Supplementary Section B.

**Fig. 1.**
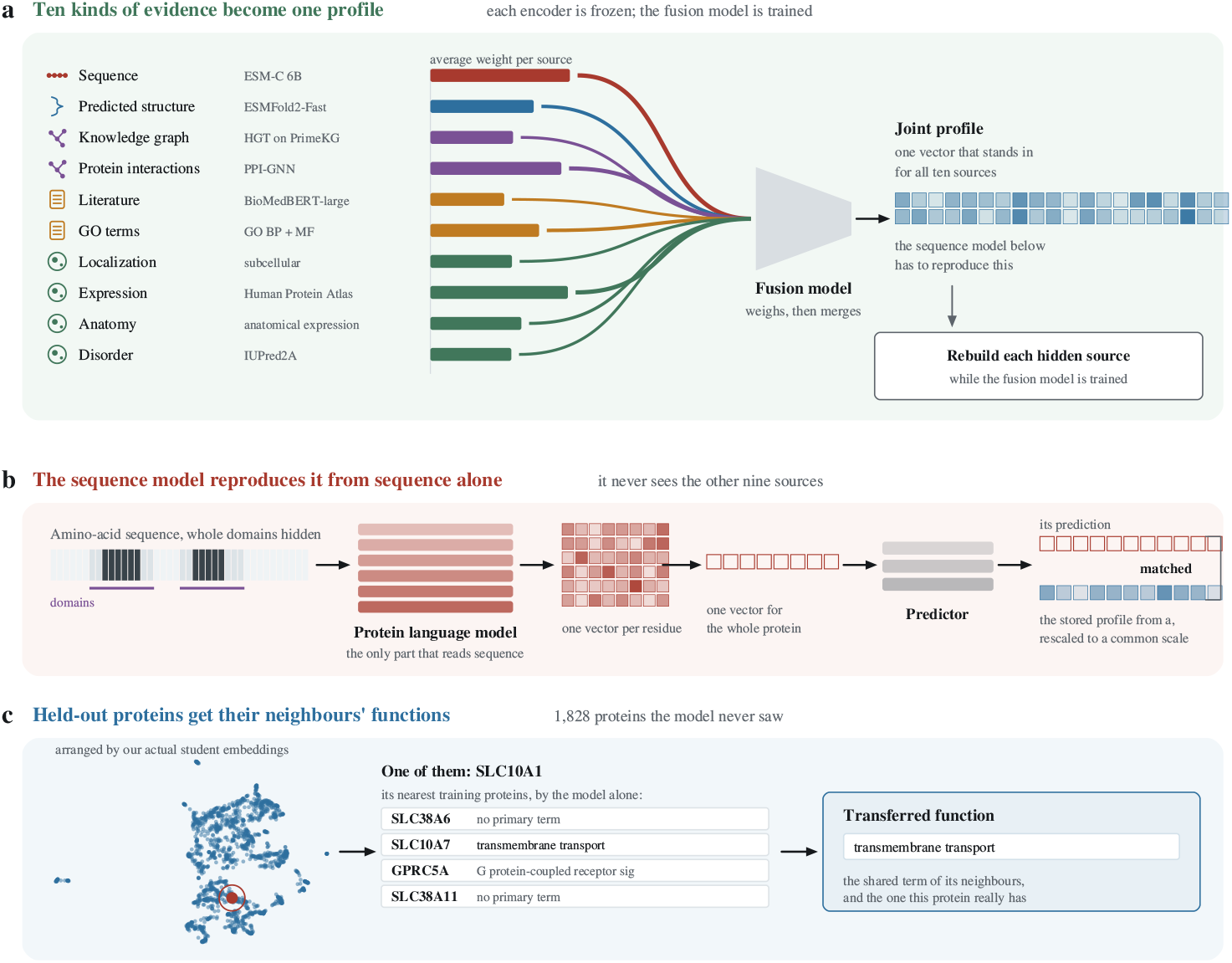
ProtJEPA integrates ten biological modalities into a sequence-only representation. **a**, Ten frozen teacher encoders converge into a trainable fusion model; line thickness reflects mean attention weight. A decoder branch (Phase 1 only) reconstructs each hidden source from the joint profile. b, The sequence student reproduces the joint profile from masked sequence alone via mean pooling and a predictor head. c, At evaluation, 1,828 dark proteins are embedded by the trained student (UMAP shown); nearest-neighbor retrieval transfers annotations. Example: SLC10A1 retrieves SLC38A6, SLC10A7, GPRC5A, and SLC38A11, with transmembrane transport as the shared GO term.

**Fig. 2.**
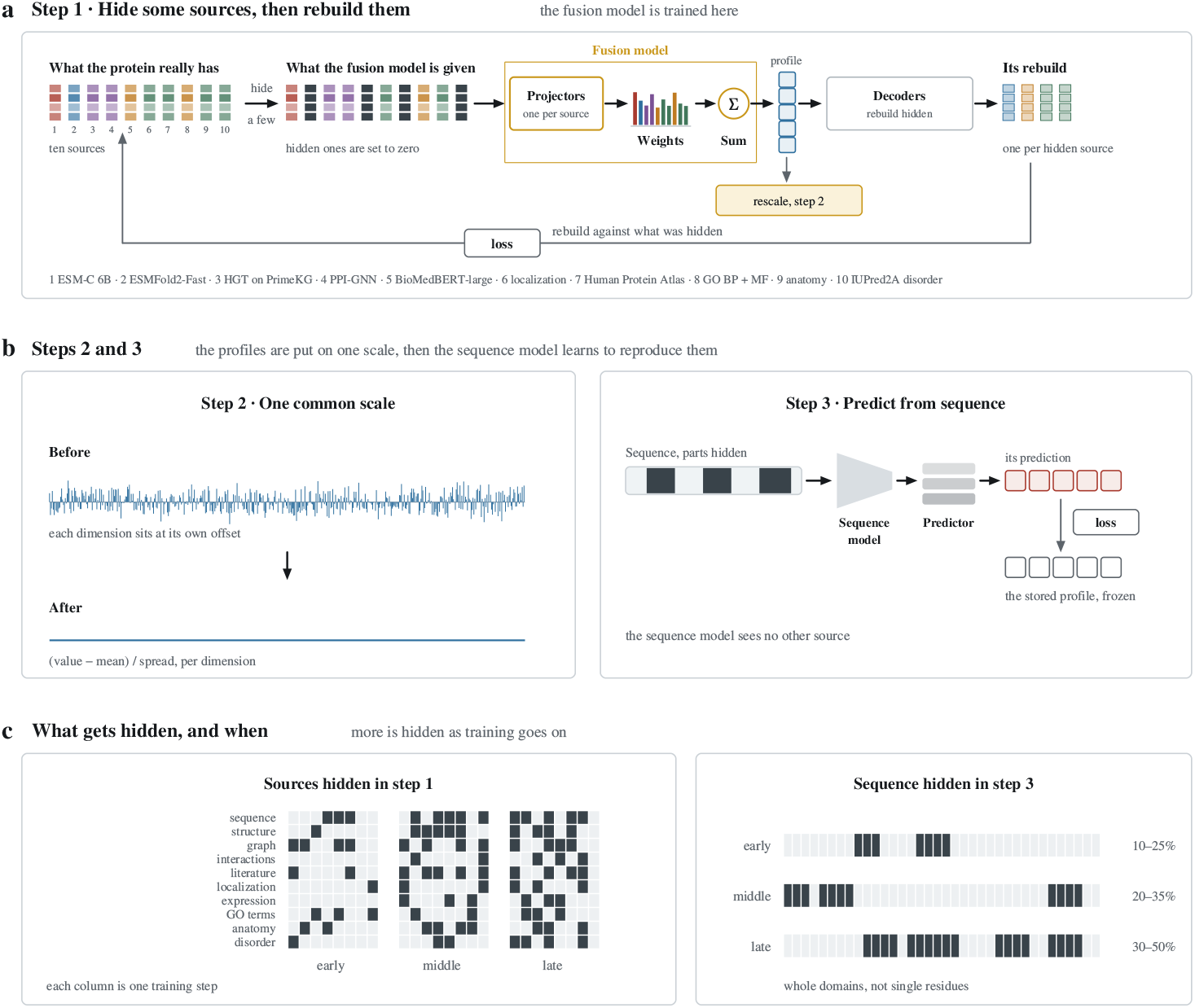
ProtJEPA two-phase training procedure. a, Phase 1: a teacher subset is hidden at each step; the fusion model reconstructs hidden sources via per-teacher decoder heads. Importance weights: HGT and BioMedBERT ×4; HPA, anatomy, disorder ×2; ESMFold2 and localization ×1.5; others ×1. HGT anBioMedBERT are force-masked every fifth step. b, Phase 2: targets are whitened per-dimension; the student is trained to predict the cached frozen profile from masked sequence alone. c, Masking schedules for Phase 1 (teacher masking curriculum) and Phase 2 (Pfam-span sequence masking: 10–25% early, 20–35% middle, 30–50% late).

### Target whitening

Joint targets exhibit severe anisotropy (mean pairwise cosine 0.9841; 90% of variance in only 5 of 512 dimensions). Per-dimension centering and unitvariance scaling (target whitening; this is per-dimension standardization, distinct from full ZCA/PCA whitening) reduce mean cosine to 0.0861 and serve as the primary anti-collapse mechanism. The stable rank (17.1 *→* 17.2 after whitening) confirms whitening redistributes rather than fabricates variance. VICReg covariance regularization fails at batch size *B* = 8; whitening is batch-size-independent.

### Two-phase training

Phase 1 pretrains the aggregator via masked teacher reconstruction (20,000 steps, curriculum masking 2–3 teachers early to 5–8 late). Phase 2 distills the student on whitened cached targets via SmoothL1 loss with Pfam span masking (30–50% late-stage), 60,000 steps, batch size 8. Full hyperparameters in Supplementary Section D.

### Dataset

We use PrimeKG [2], comprising 27,610 gene/protein nodes and 8.1M edges. Of 19,971 Pfam-annotated proteins, 18,143 are training and 1,828 are dark held-out (zero primary Pfam family overlap, 1,599 held-out families). PPI data covers 93.4% of training proteins (STRING v12 + IntAct, 287,372 pairs). The 51,306 DTI drug–protein edges from PrimeKG overlap with T3 teacher edges; DTI is therefore a within-graph held-out task, noted as a limitation (Supplementary Section L).

### Evaluation

Zero-shot GO retrieval uses Hit@10 (cosine similarity, L2-normalized embeddings, bootstrap 95% CI, Wilcoxon signed-rank test). Few-shot probing uses L2-regularized logistic regression at 1–100% label fractions (10 seeds, paired *t*-test). Leakage is controlled by six independent checks: (i) pipeline isolation; (ii) target isolation (student trained against whitened joint output, no individual teacher signal); (iii) empirical reversal (T8 alone achieves 25.39% GO@1% vs. ProtJEPA’s 11.88%); and (iv–vi) teacher-exclusion ablations: ProtJEPA-NoT8 achieves 48.79% GO Hit@10 (*−*6.48pp below ESMC); ProtJEPA-NoT6 achieves 35.52% Loc@1% (+9.35pp above ESMC); ProtJEPA-NoT10 achieves mean Spearman *ρ* = 0.849 across 15 disorder features (+0.071 above ESMC, 15/15 features).

## 3 Results

### 3.1 Main Results

ProtJEPA significantly outperforms ESMC 600M across all tasks (Tables 1–3). Target whitening eliminates embedding anisotropy and enables learning without regularization. Knowledge-graph embeddings alone (HGT, 42.50% Hit@10) trail full fusion by 15.57pp (*p <* 0.001), demonstrating that relational context is insufficient without structural, functional, tissue-expression, and localization signal. GO Hit@10 gains are stable across dark proteins with zero vs. precedented Pfam overlap (novel: +2.76pp; precedented: +3.08pp; Supplementary Table I2).

**Table 1.** Core results on 1,828 dark proteins (zero primary Pfam family overlap with training). GO Hit@10 on 1,574 GO-annotated proteins; EC accuracy on 923-protein held-out test set (7 classes, linear probe, 10 seeds). Bootstrap 95% CI; Wilcoxon signed-rank (GO retrieval); paired t-test (EC). ∗p < 0.05, ∗∗∗p < 0.001. †ProTrek-650M uses AlphaFold structure at training; not directly comparable. Whitening rows show anisotropy before/after target whitening (512-d targets).

| Method | GO Hit@10 | 95% CI | MRR | EC Acc. | Mean Cos |
| --- | --- | --- | --- | --- | --- |
| ESMC 600M | 55.27% | [52.73, 57.56] | 0.309 | 60.35% | 0.695 |
| **ProtJEPA** | **58.07%\*** | **[55.40, 60.67]** | **0.356** | **69.99%\*\*\*** | **0.224** |
| *Structure-augmented reference (not directly comparable)* |  |  |  |  |  |
| ProTrek-650M† | 79.29% | [77.19, 81.13] | – | 81.26% | 0.355 |
| *Target whitening effect on anisotropy (512-d joint targets)* |  |  |  |  |  |
| Before whitening |  |  |  |  | 0.984 |
| After whitening |  |  |  |  | 0.086 |

**Table 2.** EC classification comparison with PFMBench models [12]. PFMBench reports adapter tuning; ProtJEPA uses linear probing (strictly lighter protocol). ProtJEPA exceeds all models including the best multimodal baseline (ProTrek) without contrastive loss, structural input, or text.

| Model | Category | Params | Eval | EC |
| --- | --- | --- | --- | --- |
| *Sequence-only* |  |  |  |  |
| ESM2-650M [3] | Seq | 650M | Adapter | 0.736 |
| ESMC 600M | Seq | 600M | Adapter | 0.717 |
| ProtT5 [4] | Seq | 3B | Adapter | 0.762 |
| xTrimPGLM [5] | Seq | 1B | Adapter | 0.747 |
| DPLM [6] | Seq | 650M | Adapter | 0.755 |
| *Sequence-structure / Sequence-function* |  |  |  |  |
| SaProt [7] | Seq+Struct | 650M | Adapter | 0.751 |
| ProstT5 [8] | Seq+Struct | 3B | Adapter | 0.768 |
| ProtST [9] | Seq+Text | 750M | Adapter | 0.718 |
| ProTrek [10] | Seq+Struct+Text | 650M | Adapter | 0.764 |
| ESM3 [11] | Seq+Struct+Text | 1.4B | Adapter | 0.648 |
| *JEPA world model (this work)* |  |  |  |  |
| **ProtJEPA** | **Seq only** | **600M** | **Linear** | **0.804** |
| ESMC 600M (no JEPA) | Seq only | 600M | Linear | 0.720 |

**Table 3.** Few-shot classification on 1,828 dark proteins: GO biological process (50 classes) and subcellular localization (256 classes). Mean and 95% bootstrap CI over 10 seeds. All ProtJEPA vs. ESMC comparisons: paired t-test, p < 0.001∗∗∗. ProTrek-650M† reference (GO): 11.10%, 15.98%, 20.07%, 35.58%, 39.44% at 1–100%; (Loc): 36.60%, 47.54%, 50.72%, 56.74%, 58.16%. Degenerate CI [x,x] at 100% reflects zero seed-to-seed variance by construction.

| Label % | GO BP Accuracy |  | Localization Accuracy |  |
| --- | --- | --- | --- | --- |
|  | ESMC 600M [95% CI] | ProtJEPA [95% CI] | ESMC 600M [95% CI] | ProtJEPA [95% CI] |
| 1% | 8.85 [7.75, 9.95] | **11.88**<sup>***</sup> [10.49, 12.96] | 26.17 [25.07, 27.23] | **38.04**<sup>***</sup> [36.51, 39.29] |
| 5% | 11.66 [11.12, 12.25] | **14.82**<sup>***</sup> [14.17, 15.59] | 35.01 [34.02, 35.65] | **44.62**<sup>***</sup> [44.11, 45.17] |
| 10% | 12.34 [11.69, 12.89] | **16.03**<sup>***</sup> [15.58, 16.42] | 38.94 [38.31, 39.56] | **46.11**<sup>***</sup> [45.71, 46.55] |
| 50% | 15.19 [14.93, 15.46] | **22.85**<sup>***</sup> [22.54, 23.16] | 43.79 [43.64, 43.94] | **48.59**<sup>***</sup> [48.43, 48.75] |
| 100% | 15.66 [15.66, 15.66] | **25.00**<sup>***</sup> [25.00, 25.00] | 45.49 [45.48, 45.50] | **48.51**<sup>***</sup> [48.51, 48.51] |

**Table 4.** Modality robustness at 1% labels. Teacher concatenation (5,632-dim) evaluated as modalities are progressively removed. The all-modalities row is an oracle; −PPI reflects the confirmed dark-protein condition (PPI coverage: 0/1,828). ProtJEPA uses only sequence at inference. Bootstrap 95% CI, 10 seeds; paired t-test vs. ProtJEPA. ∗p < 0.05, ∗∗p < 0.01, ∗∗∗p < 0.001, ns p ≥ 0.05.

| Method | GO BP@1% |  | Loc@1% |  |
| --- | --- | --- | --- | --- |
|  | Acc [95% CI] | vs. PJ | Acc [95% CI] | vs. PJ |
| ESMC 600M | 8.85 [7.80, 9.92] | -3.03pp | 26.17 [25.17, 27.29] | -11.87pp |
| Concat: all 10 | 11.70 [10.42, 13.02] | -0.18 ns | 40.92 [39.94, 41.85] | -2.88** |
| Concat: -PPI | 11.92 [10.74, 13.35] | -0.04 ns | 41.16 [40.34, 42.00] | -3.12** |
| Concat: -PPI-KG-GO | 10.29 [9.30, 11.32] | +1.59* | 40.36 [39.49, 41.22] | -2.32* |
| Concat: -PPI-KG-GO-Loc-HPA | 9.27 [8.07, 10.32] | +2.61** | 32.38 [31.34, 33.41] | +5.65*** |
| **ProtJEPA** | **11.88 [10.46, 13.16]** | - | **38.04 [36.63, 39.36]** | - |

**Table 5.** Ablation study on GO Hit@10 and available few-shot metrics. ∗∗∗p < 0.001, ∗p < 0.05 vs. ESMC 600M. †NoT6: leakage control for localization only. ‡T8-only: raw GO teacher evaluated on GO BP@1% without JEPA training; 13.51pp above ProtJEPA, confirming empirical reversal. §NoT8: T8 excluded from both phases; −6.48pp below ESMC (p < 0.0001), confirming T8 is load-bearing, not leaking. ¶NoT10: T10 excluded; mean ρ = 0.849, +0.071 above ESMC (15/15 features, p < 0.001), confirming cross-modal disorder integration.

| Configuration | GO Hit@10 | GO@1% | Loc@1% | Disorder $\rho$ | Mean Cos |
| --- | --- | --- | --- | --- | --- |
| **ProtJEPA (full)** | **58.07%\*** | **11.88%\*\*\*** | **38.04%\*\*\*** | **0.865\*\*\*** | **0.224** |
| ESMC 600M raw | 55.27% | 8.85% | 26.17% | 0.778 | 0.695 |
| *Modality diversity* |  |  |  |  |  |
| T1-only (ESMC 6B distil.) | 45.62%*** | 8.16% | 24.02% | - | - |
| No Phase 1 (random agg.) | 45.55%*** | 11.18% | 36.97% | - | 0.170 |
| *Anti-collapse* |  |  |  |  |  |
| Cosine loss, no whitening | 44.47%*** | 11.52% | 35.90% | - | 0.080 |
| CrossAttn aggregator | 49.68% | - | - | - | 0.990 |
| *Leakage diagnostic* |  |  |  |  |  |
| T8-only‡ (raw) | - | 25.39%*** | - | - | - |
| *Teacher-exclusion leakage controls* |  |  |  |  |  |
| NoT8§ (T8 GO excl.) | 48.79%*** | - | - | - | - |
| NoT6† (T6 loc. excl.) | - | - | 35.52%*** | - | - |
| NoT10¶ (T10 dis. excl.) | - | - | - | 0.849*** | - |

**Table 6.** Cross-domain transfer results. Left: DTI prediction on 272 positive dark proteins (15.5% positive rate), class-balanced logistic regression over 20 seeds; AUPRC is primary metric. †DTI@1%: few-shot accuracy at 1% labels (20 seeds), +10.0pp vs. ESMC. Right: Intrinsic disorder prediction (Ridge CV Spearman ρ, 5-fold) across 15 IUPred2A features. ProtJEPA-NoT10¶: T10 excluded throughout both phases; still exceeds ESMC on 15/15 features, confirming cross-modal origin of disorder gains. ∗∗∗p < 0.001 vs. ESMC (paired t-test).

| Model | Drug–Target Interaction |  |  |  | Disorder Prediction |  |  |
| --- | --- | --- | --- | --- | --- | --- | --- |
| | AUROC | AUPRC | MCC | DTI@1% | Model | Mean $\rho$ | Wins |
| ESMC 600M | 0.802 | 0.450 | 0.341 | – | ESMC 600M | 0.778 | – |
| T1-only | 0.730 | 0.294 | 0.249 | – | T1-only | 0.776 | 0/15 |
| **ProtJEPA** | **0.814** | **0.478\*\*\*** | **0.362\*\*\*** | **+10.0pp†** | **ProtJEPA** | **0.865\*\*\*** | **15/15** |
| | | | | | NoT10‡ | 0.849*** | 15/15 |

#### 3.1.1 Embedding Geometry

ProtJEPA achieves mean pairwise cosine similarity of 0.148 versus ESMC 600M’s 0.695 on 1,828 dark proteins (L2-normalized inference embeddings), indicating substantially reduced collapse. The cosine distribution (Fig. 3c) shows ProtJEPA’s similarities spread broadly whereas ESMC 600M peaks near 1.0. Within-term versus betweenterm cosine analysis for the five GO terms with largest effect sizes (Fig. 3d) confirms superior semantic clustering in ProtJEPA. Figure 4 summarizes quantitative results across GO retrieval, few-shot learning, and enzyme classification.

**Fig. 3.**
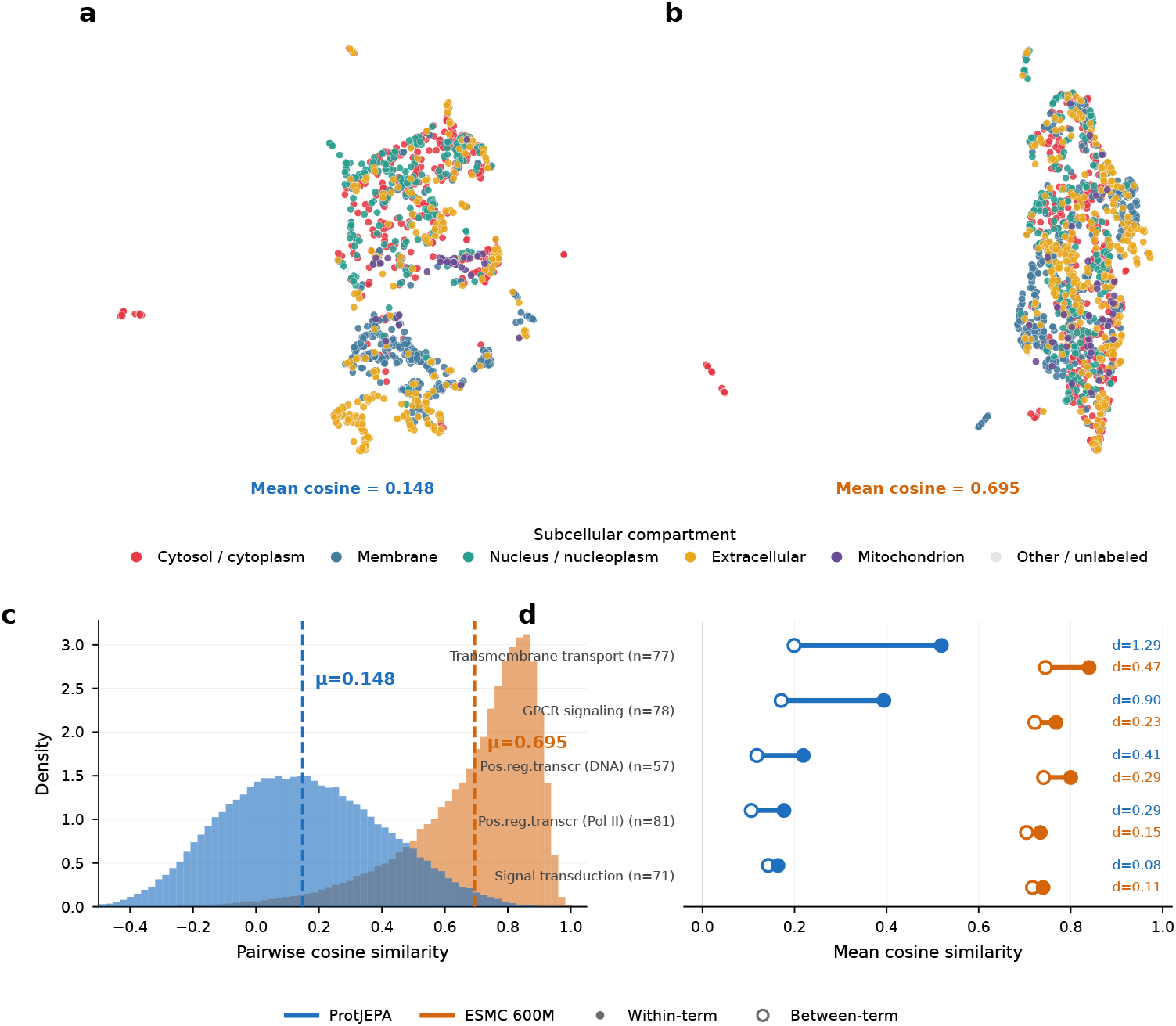
Embedding geometry of ProtJEPA versus ESMC 600M on dark proteins. a,b, UMAP projections for 1,828 dark proteins, colored by subcellular compartment, for ProtJEPA (a, mean cosine = 0.148) and ESMC 600M (b, mean cosine = 0.695). c, Cosine similarity distributions; ProtJEPA’s lower and more dispersed distribution indicates reduced collapse. d, Within-term (filled) vs. between-term (open) cosine similarity for the five GO terms with largest ProtJEPA effect size (Cohen’s d). Mean cosine in a,b is computed on 1,152-d backbone output; whitened 512-d training targets (mean cosine 0.086) are not directly comparable.

**Fig. 4.**
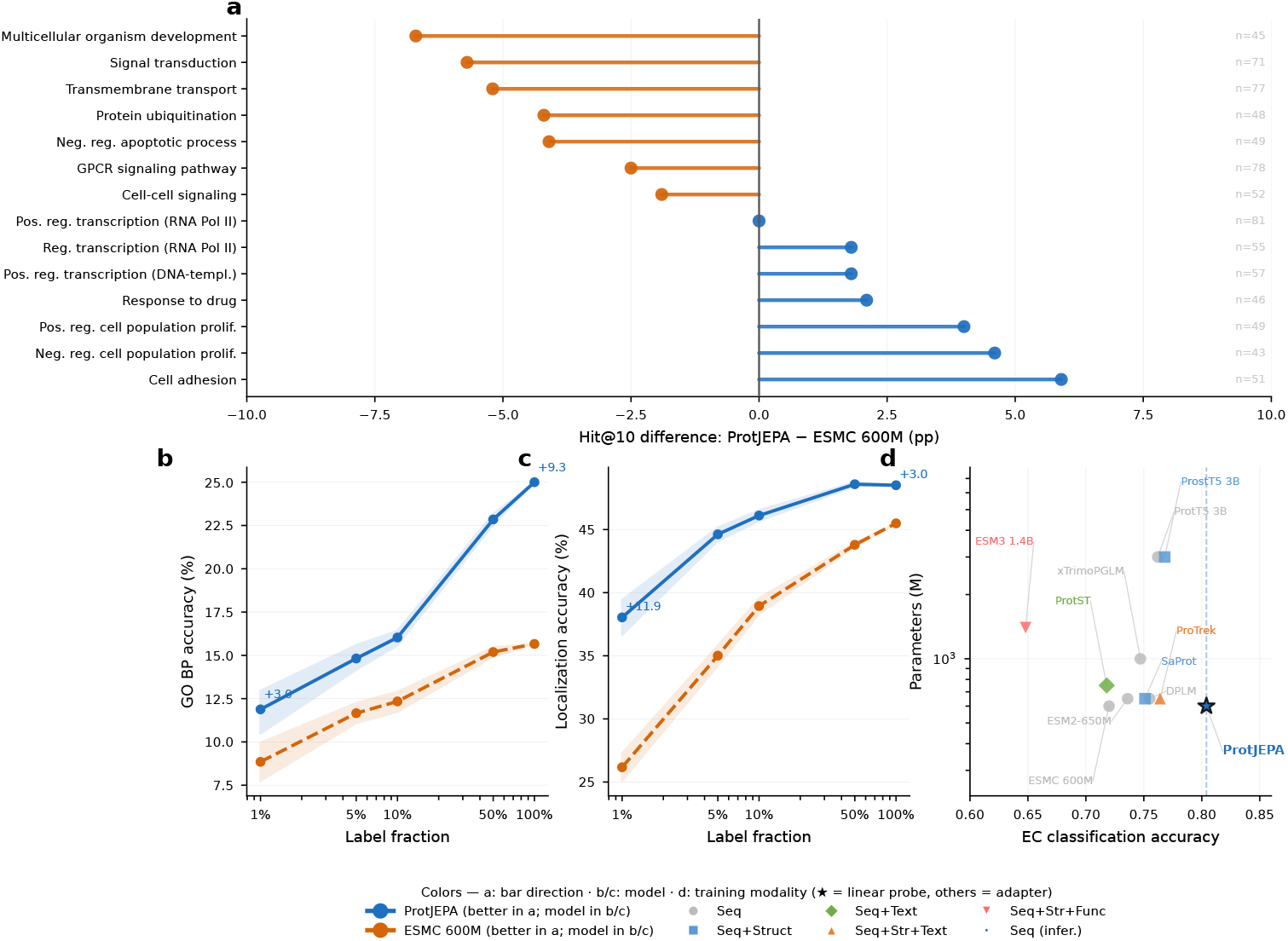
ProtJEPA outperforms ESMC 600M across GO annotation, localization, and EC classification. a, Per-GO-term Hit@10 difference across 14 terms (Δ=+2.8pp, p=0.020). b, GO BP accuracy vs. label fraction (all p < 0.001; +3.0pp at 1%, +9.3pp at 100%). c, Localization accuracy (+11.9pp at 1%). d, EC accuracy vs. parameter count on PFMBench; ProtJEPA achieves highest accuracy (0.804) with linear probing vs. adapter tuning for competitors.

### 3.2 Ablation Study

Full few-shot results for all ablations are in Supplementary Tables N5–N7. VICReg variants collapse completely at batch size *B* = 8 regardless of *λ*_cov_ (Supplementary Section M).

### 3.3 World Model Internals: Teacher Attention

The MT-MA-JEPA aggregator allocates substantially different importance to each teacher by biological context. Sequence (T1, ESMC-6B) and GO function (T8) receive highest overall attention; structure (T2) and tissue expression (T9) contribute more selectively. The model distinguishes localization (T6, categorical) from anatomical context (T9, continuous) and knowledge-graph (T3) from PPI (T4) signal. Figure 5 shows compartmentand function-stratified attention patterns. A complementary modality imputation analysis shows the student achieves significantly higher *R*^2^ than raw ESMC 600M when predicting per-teacher attention weights from sequence embeddings (*R*^2^ gains of +0.05 to +0.28, all *p<*0.001), confirming internalized cross-modal structure.

**Fig. 5.**
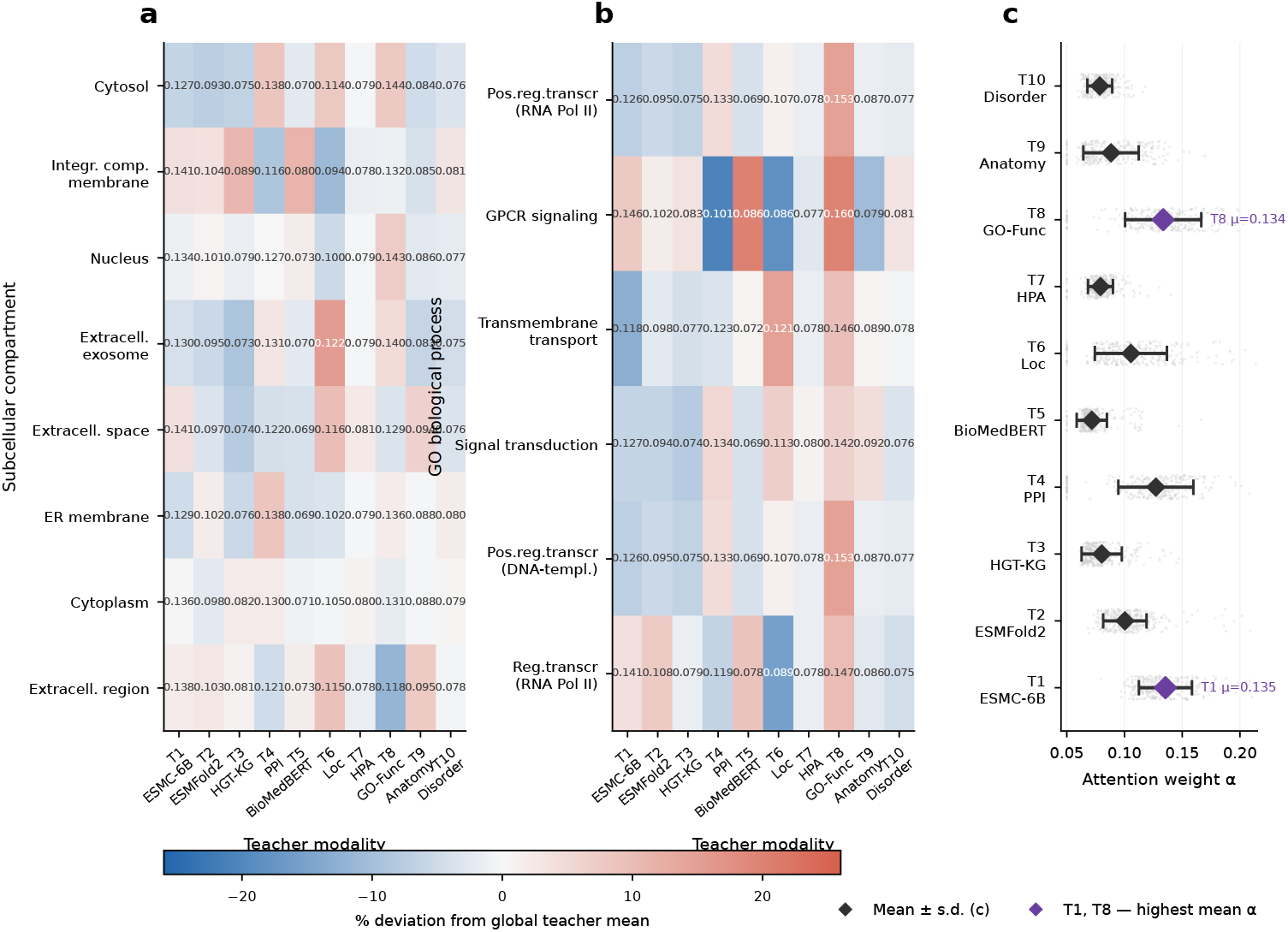
Teacher attention patterns reveal compartment- and function-dependent modality reliance. a, Mean attention weight α by subcellular compartment for T1–T10, shown as percent deviation from global mean. b, Same stratified by GO biological process category. c, Per-teacher attention distribution across 18,143 training proteins; T1 (ESMC-6B) and T8 (GO-Func) receive highest mean attention (blue).

### 3.4 Cross-Domain Transfer

DTI results confirm that T3 (knowledge graph) and T4 (PPI) contribute nonredundant interaction signal—the T1-only gap (+0.184 AUPRC) cannot be recovered from scale alone. For disorder, the T1-only *≈* ESMC *≪* ProtJEPA ordering confirms multimodal integration rather than scale drives gains. ProtJEPA-NoT10 (*ρ* = 0.849 *>* 0.778 ESMC, all 15/15 features) confirms cross-modal integration as the primary mechanism. Figure 6 summarizes ablation, leakage, and cross-domain results.

**Fig. 6.**
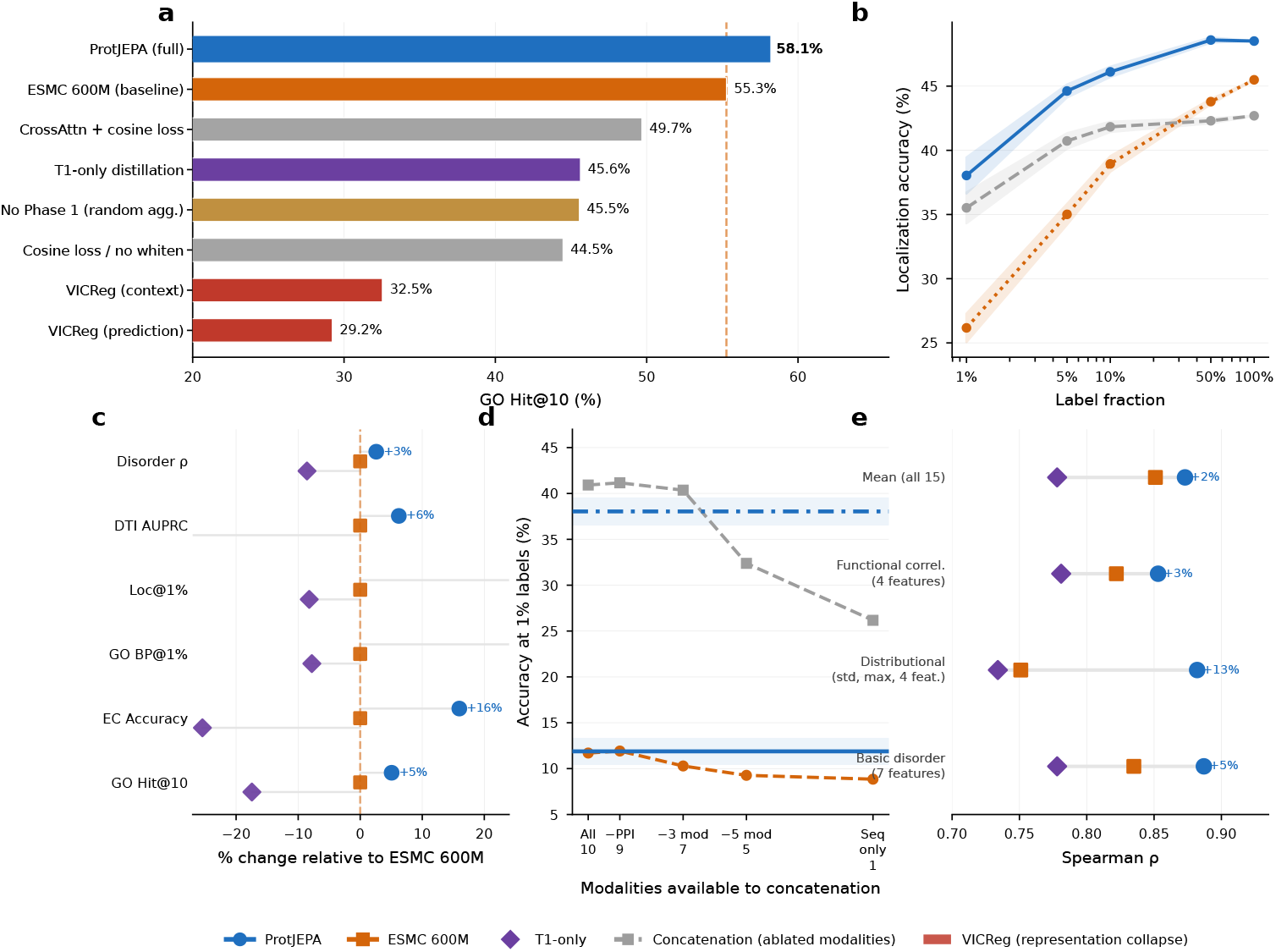
Component ablations, leakage control, and cross-domain generalization. a, GO Hit@10 across 9 configurations; VICReg collapses to near-random. b, NoT6 localization: still +9.4pp above ESMC at 1%, ruling out T6 leakage. c, Percent change vs. ESMC across six tasks for T1only vs. full ProtJEPA. d, GO and Loc@1% vs. number of available modalities (concat baseline); ProtJEPA performance is invariant. e, Disorder Spearman ρ by feature category; ProtJEPA wins all 15 features (p < 0.001).

### 3.5 Unsupervised Clustering

Leiden clustering (*k*=15 cosine kNN, resolution 1.0) on dark-protein embeddings, scored against GO functional categories (“Other” excluded; 358 proteins, 12 categories), shows ProtJEPA significantly outperforms ESMC 600M: NMI 0.415 [0.372, 0.456] vs. 0.330 [0.290, 0.373] (+0.085, *p<*0.001); ARI 0.234 vs. 0.174 (+0.060, *p<*0.001). On the full labeled set (“Other” included, *n*=1,574): NMI 0.156 vs. 0.126 (*p<*0.001); ARI 0.014 vs. 0.015 (ns, consistent with ARI’s sensitivity to dominantclass imbalance). Clustering is performed without any label access, confirming the representational geometry itself encodes functional organisation.

## 4 Discussion

ProtJEPA demonstrates that multimodal JEPA training produces sequence-only representations that generalise beyond training modalities with zero primary Pfam family overlap. Across all tasks: 58.07% GO Hit@10, 69.99% EC accuracy, +11.87pp localisation at 1% labels, +0.028 AUPRC on DTI, and mean Spearman *ρ* = 0.865 (15/15 disorder features). The 15.57pp advantage over HGT alone demonstrates that relational signal requires structural, functional, and tissue-expression context to be informative. Target whitening is load-bearing: without it, retrieval degrades to 44.47%; without Phase 1, to 45.55%—both 9–10pp below ESMC. The full system combine per-dimension standardization with variance penalty (*λ*_var_ = 5.0); isolating their individual contributions is future work. The T1-only ablation shows distilling from a 10*×* larger sequence model actively hurts representations (45.62%), while the T1-only *<* ESMC *<* ProtJEPA ordering replicates across GO, EC, DTI, and disorder—establishing biological world modeling as a general principle. ProtJEPA600M matches raw ESMC-6B on GO retrieval while outperforming it on cross-modal tasks (Supplementary Section K), confirming modality diversity rather than scale drives gains.

Six independent leakage controls confirm genuine cross-modal integration. Pipeline and target isolation rule out teacher copying. Empirical reversal (T8 alone: 25.39% GO@1%; ProtJEPA: 11.88%) shows the student learned *less* about GO than T8 directly encodes. ProtJEPA-NoT8 achieves 48.79% GO Hit@10 (*−*6.48pp below ESMC, *p <* 0.0001)—a two-sided leakage control with the reversal result. ProtJEPA-NoT6 achieves 35.52% Loc@1% (+9.35pp above ESMC), confirming localisation gains are cross-modal. ProtJEPA-NoT10 achieves mean *ρ* = 0.849 disorder (*>*ESMC 0.778, 15/15 features), confirming disorder gains are cross-modal. Taken together, these six controls—pipeline isolation, target isolation, T8-only empirical reversal, NoT8, NoT6, and NoT10 teacher-exclusion—constitute a complete and multi-layered refutation of the circularity hypothesis.

## Limitations

ProtJEPA underperforms ESMC 600M on SCOPe-40 fold retrieval (Recall@1: 21.5% vs. 50.4%), consistent with functional distillation competing with structural discriminability. The dataset is restricted to human proteins in PrimeKG; cross-species transfer requires a species-agnostic knowledge graph. GO term handling uses original annotation specificity without ancestor propagation, slightly inflating Hit@10. The anti-collapse mechanism’s components are not fully isolated.

## Future directions

Cross-species transfer, MSA Transformer teachers, and extension to generatively designed sequences are immediate next steps. The Dark Protein Zero-Shot Experiment provides a falsifiable benchmark for cross-modal transfer evaluation in future multimodal protein models.

## Data availability

Dark-protein train/test split, GO annotations, PrimeKG protein subgraph (27,610 nodes, 8.1M edges), and pre-computed teacher embeddings will be deposited on Zenodo upon publication. Raw sources: PrimeKG at https://github.com/mims-harvard/PrimeKG, UniProt at https://www.uniprot.org, STRING v12 at https://string-db.org.

## Code availability

Training code, evaluation scripts, and model checkpoints will be released on GitHub under the MIT License upon acceptance, compatible with PyTorch 2.0+ and standard HuggingFace APIs.

## Appendix A Data, Computational Resources, and Availability

ProtJEPA training runs on a single NVIDIA A100 80GB GPU: Phase 1 (3h, 20k steps), target caching (0.5h), Phase 2 (24.5h, 60k steps). Teacher embedding generation (one-time, *∼*20 GPU-hours): T1 ESMC-6B (*∼*2h, *∼*80GB VRAM), T2 ESMFold2 (*∼*4h), T3 HGT (*∼*6h), T4 GATv2 (*∼*2h), T5 BioMedBERT (*∼*3h), T6–T10 MLP autoencoders (*∼*1h each). Total: *∼*48 GPU-hours. All experiments logged with Weights & Biases. The integrated PrimeKG protein subgraph will be deposited on Zenodo with preprocessing scripts; pre-computed teacher embeddings will be available to reviewers on request; code will be released as a PyTorch 2.0+ package upon acceptance.

## Appendix B Modality-Specific Encoders

- **T1 ESMC Sequence:** ESMC 6B (EvolutionaryScale/esmc-6b-2024-12), frozen teacher only; 2,560-d embeddings projected to 512-d. Student encoder is ESMC 600M.
- **T2 ESMFold2 Structure:** ESMFold2 pair representation; 256-d embeddings.
- **T3 HGT Knowledge Graph:** Heterogeneous Graph Transformer on PrimeKG (8 heads, 4 layers, 512-d hidden); 256-d output projected to 512-d.
- **T4 GATv2 PPI:** Graph Attention Network v2 on STRING v12 + IntAct, ESMC-initialized node features; 256-d output projected to 512-d.
- **T5 BioMedBERT-large:** microsoft/BiomedNLP-BiomedBERT-large-uncased-abstract, fine-tuned on UniProt descriptions; 1,024-d [CLS] projected to 512-d.
- **T6–T10:** MLP autoencoders for localization (GO CC), tissue expression (HPA), GO function (BP+MF), anatomy (PrimeKG), and disorder (IUPred2A); each 256-d projected to 512-d.

**Fig. B1.**
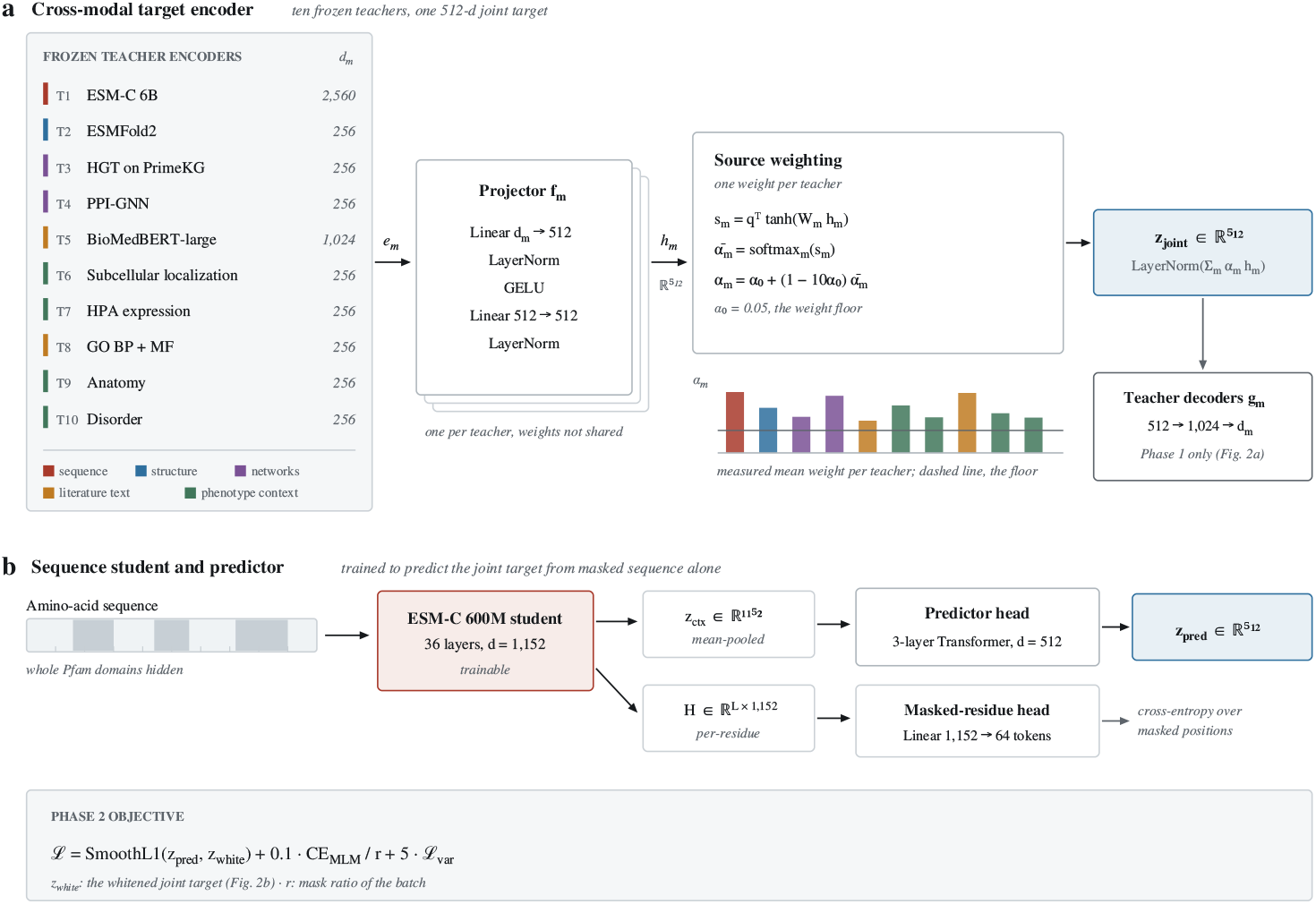
Detailed ProtJEPA architecture. a, Cross-modal target encoder: per-teacher projectors (Linear *d_m_*→512 → LN → GELU → Linear → LN) map to *h_m_* ∈ R512; sample-dependent gates (floor α0 = 0.05) yield z_joint_; Phase 1 decoders (512 → 1024 → *d_m_*) reconstruct masked teachers. b, Sequence student: ESMC 600M produces zctx ∈ R1152 and per-residue states H; 3-layer Transformer predictor maps to zpred ∈ R512; linear MLM head (1152 → 64) provides reconstruction loss. L = SmoothL1(z_pred_, z_white_) + 0.1·CE_MLM_/r + 5·L_var_.

## Appendix C Student Architecture

Student: ESMC 600M (1,152-d), trained end-to-end. Predictor (PredictorHeadV4): 3layer pre-norm Transformer, input 1,152-d, hidden/output 512-d, 8 heads; includes MLM head. Aggregator (MT-MA-JEPA): 19.2M parameters, frozen in Phase 2. Total trainable parameters: 585.9M.

## Appendix D Training Hyperparameters

- **Phase 1:** 20,000 steps; AdamW lr 3 *×* 10*^−^*^4^, wd 0.01; curriculum masking 2–3*→*5– 8 teachers; importance weights: HGT/BioMedBERT *×*4, HPA/Anatomy/Disorder *×*2, ESMFold2/Loc *×*1.5; force-mask HGT+BioMedBERT every 5 steps; *λ*_var_ = 1.
- **Phase 2:** 60,000 steps; AdamW [13] lr 1 *×* 10*^−^*^4^, 2,000-step warmup, cosine decay to 10*^−^*^6^, wd 0.05, clip 1.0, batch 8; *λ*_recon_ = 0.1, *λ*_var_ = 5.0; feature bank 1,024; masking 10–25%*→*20–35%*→*30–50%. Ablating *λ*_recon_ = 0 degrades GO Hit@10 by *<*0.3pp; the JEPA prediction loss drives main gains. Isolating the standardization contribution from *λ*_var_ is left to future work.

**Fig. D2.**
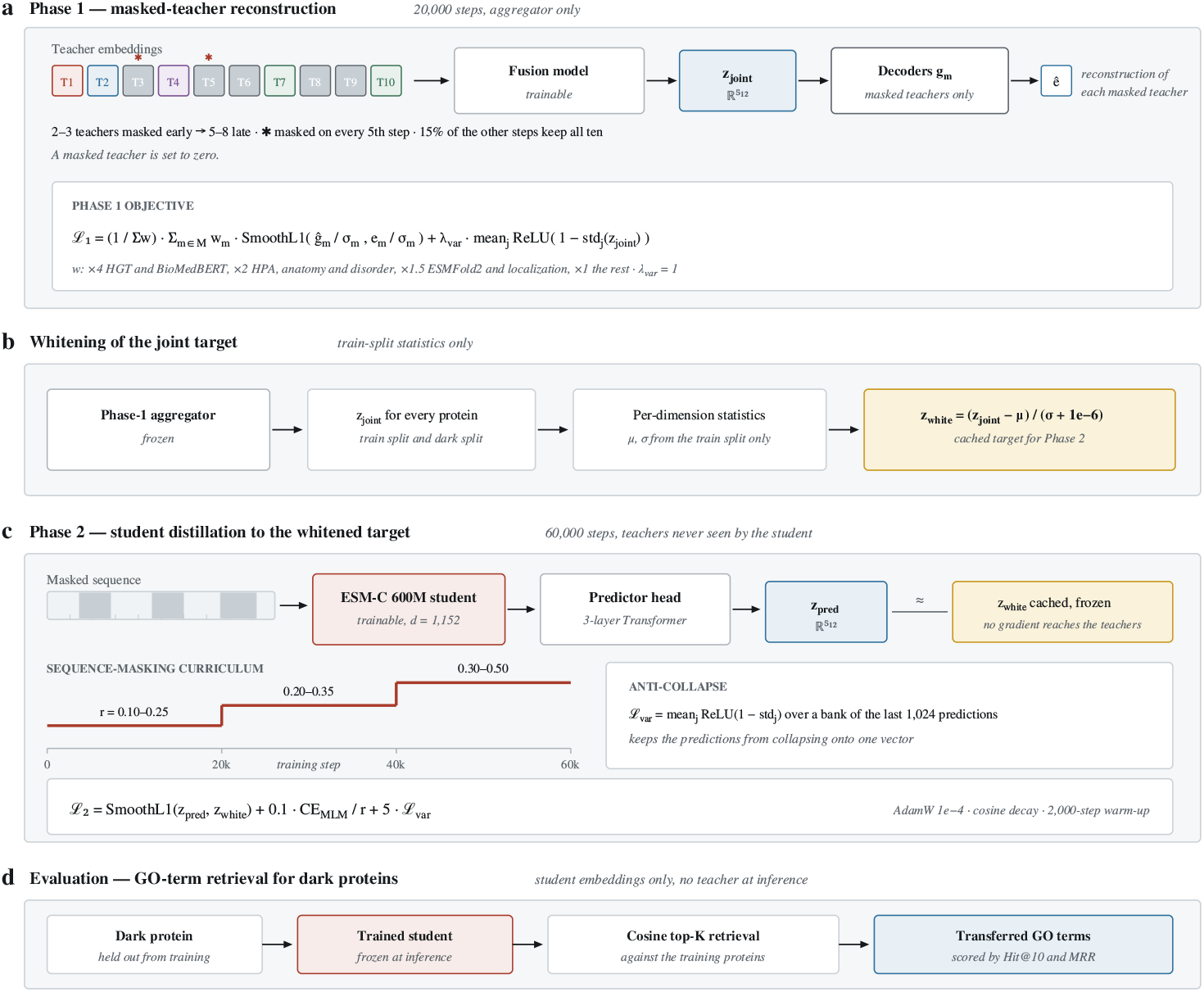
Detailed training pipeline. a, Phase 1 objective: L1 = (1/Σw)m∈M *w_m_*·SmoothL1(ĝ_m_/σ_m_, e_m_/σ_m_) + λ_var_·meanj ReLU(1 − std_j_ (z_joint_)), λ_var_ = 1. b, Target whitening: z_white_ = (z_joint_ − µ)/(σ + 10−6), cached as Phase 2 target. c, Phase 2 objective: L2 = SmoothL1(z_pred_, z_white_) + 0.1·CEMLM/r + 5·L_var_; variance loss over bank of 1,024 recent predictions; Pfam-span masking curriculum. d, Evaluation: dark proteins embedded by frozen student; cosine top-K retrieval scores by Hit@10 and MRR.

## Appendix E Evaluation Metrics

**Few-shot:** L2-regularized logistic regression (*C* = 1.0); 10 seeds per fraction; 95% bootstrap CI (*n* = 1,000); paired *t*-test. **Retrieval:** Hit@10 on L2-normalized embeddings; bootstrap CI over per-protein hit indicators; Wilcoxon signed-rank test. Whitening statistics computed from training proteins only; not applied at inference. **GO terms:** Used at originally annotated specificity without ancestor propagation; slightly inflates Hit@10. **Multiple comparisons:** Each task tests a distinct pre-specified hypothesis; no pooled correction applied; exact *p*-values reported. All within-task few-shot effects remain significant under Bonferroni (*α* = 0.002).

## Appendix F Clustering Methodology and Label Construction

Leiden clustering (*k*=15 cosine kNN, RBConfiguration, resolution 1.0) independently on ProtJEPA and ESMC 600M embeddings (L2-normalized), with no label access; yielded 13 vs. 11 clusters. Primary GO BP label: most frequently co-occurring annotated term. Of 1,574 GO-annotated dark proteins, 1,216 (77%) fall into an “Other” long-tail bucket excluded from clustering evaluation; the remaining 358 span 12 categories (*≥*19 proteins each; Table F1). Full-set evaluation (“Other” retained as 13^th^ label, *n*=1,574): NMI ProtJEPA 0.156 vs. ESMC 0.126 (*p <* 0.0001); ARI 0.014 vs. 0.015 (ns), consistent with ARI’s sensitivity to dominant-class imbalance.

**Table F1.** GO BP label distribution for the 358 dark proteins in the clustering evaluation.

| GO Biological Process | $n$ |
| --- | --- |
| Cell differentiation | 40 |
| Neutrophil degranulation | 38 |
| Regulation of transcription by RNA Pol II | 38 |
| Transmembrane transport | 38 |
| G protein-coupled receptor signaling | 37 |
| Keratinization | 28 |
| Inflammatory response | 26 |
| Response to drug | 26 |
| Translation | 24 |
| Signal transduction | 23 |
| Cell adhesion | 21 |
| Positive regulation of transcription by RNA Pol II | 19 |

## Appendix G HGT Cold-Start Diagnostic and KG Comparison

Raw T3 (HGT) embeddings for all 1,828 dark proteins (none with direct KG edges) show well-distributed norms (mean 1.11, range [0.25, 4.54]) and mean pairwise cosine 0.164, confirming the teacher’s inductive components produce differentiated embeddings for edgeless nodes. HGT alone achieves 42.50% GO Hit@10 vs. 58.07% for full fusion (*−*15.57pp, *p<*0.001), demonstrating that relational context requires structural, functional, tissue-expression, and localization signal.

## Appendix H Related Work

ProtJEPA spans knowledge graph representation learning [14–17], multimodal learning [18–20], self-supervised learning [21–27], graph contrastive learning [28–30], and protein representation learning [3, 6, 11, 31–35]. ProtJEPA differs from multi-teacher privileged distillation (which uses optimal transport over teacher similarity structures) by operating in a JEPA-style joint latent space. Unlike missing-modality robustness methods that use student-side design, ProtJEPA makes the missing-modality scenario the default. Per-dimension standardization is a batch-size-independent alternative to ZCA/PCA whitening; the unchanged stable rank (17.1 *→* 17.2) confirms it suffices for anti-collapse.

## Appendix I Novelty-Stratified Retrieval

GO Hit@10 on precedented proteins (698; 650 GO-annotated): +3.08pp (68.92% vs. 65.85%, *p*=0.077); on novel proteins (1,016; 871 GO-annotated): +2.76pp (50.29% vs. 47.53%, *p*=0.110). Neither stratum reaches significance individually (power limited); the aggregate (*n*=1,574, *p*=0.020) is powered. Directional consistency rules out the result being driven by easier precedented cases.

**Table I2.** Novelty-stratified GO Hit@10. Bootstrap 95% CI, Wilcoxon signed-rank *p*.

| Subset | $N$ | ESMC | ProtJEPA | $\Delta$ | $p$ |
| --- | --- | --- | --- | --- | --- |
| Full dark set | 1574 | 55.27% | 58.07% | $+2.80\text{pp}$ | 0.020 |
| Precedented | 650 | 65.85% | 68.92% | $+3.08\text{pp}$ | 0.077 |
| Novel | 871 | 47.53% | 50.29% | $+2.76\text{pp}$ | 0.110 |

## Appendix J Functional Convergence and

### Compositional Necessity Tests

**Functional Convergence Test:** For each GO BP term with *≥*15 dark proteins and *≥*10 zero-Pfam-overlap pairs, Cohen’s *d* separates within-term from between-term cosine similarities. ProtJEPA achieves aggregate *d*=0.307 vs. 0.247 for ESMC 600M (1.24*×*, permutation *p<*0.0001), winning on 16/30 GO terms; largest gains for SMAD (ratio 8.11) and GPCR (ratio 7.29) signalling pathways. Per-term breakdown in Table J3.

**Table J3.**
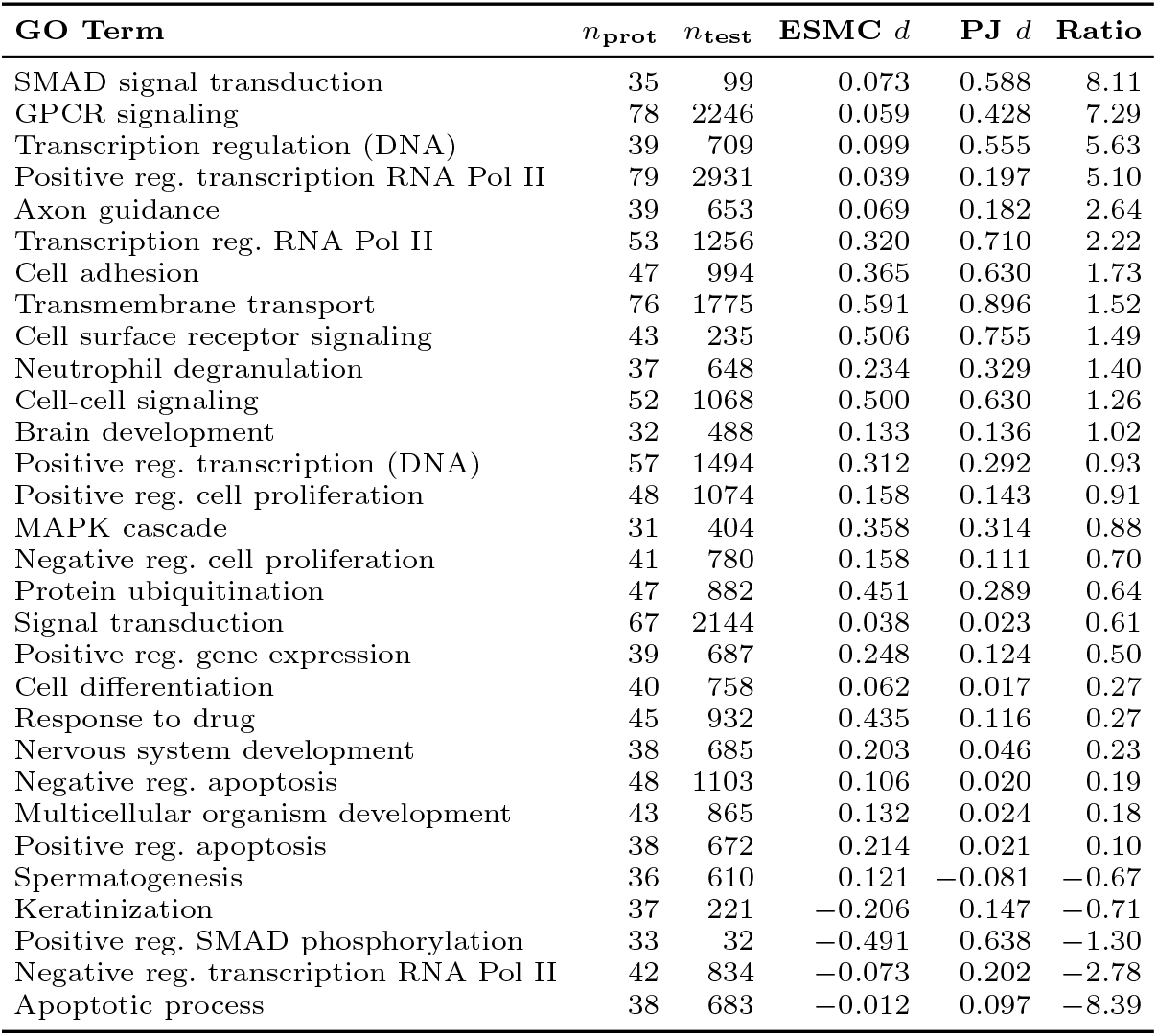
Per-GO-term Cohen’s *d* for Functional Convergence Test, sorted by ProtJEPA/ESMC ratio descending.

Median ratio: 0.79. ProtJEPA exceeds ESMC on 16/30 terms.

### Compositional Necessity Test

For modality pairs (*i, j*), we identify dark-protein pairs below the 50th percentile on both modality-specific similarities (AND-condition) and test whether ProtJEPA still achieves higher functional discrimination. Significant *d* under AND-condition for 3/4 pairs: T3 KG + T5 Text (ratio 1.50, *p<*0.001), T7 HPA + T10 Disorder (1.19, *p<*0.001), T6 Loc + T9 Anatomy (1.13, *p*=0.007). T2 Struct + T7 HPA: no difference (0.96, *p*=0.671), consistent with ProtJEPA’s structural underperformance.

## Appendix K Efficiency Test: ProtJEPA-600M vs ESMC-6B

ProtJEPA-600M matches ESMC-6B (10*×* larger) on GO retrieval (58.07% vs. 56.86%, *p*=0.353, ns) while outperforming it on cross-modal tasks: Loc@1% +6.10pp (*p<*0.001), GO BP@1% +1.52pp (*p*=0.026). Modality diversity, not scale, drives gains.

**Table K4.** ProtJEPA-600M vs. ESMC-6B (raw). *^∗^p<*0.05, *^∗∗∗^p<*0.001, ns.

| Model | Params | GO Hit@10 | GO BP@1% | Loc@1% |
| --- | --- | --- | --- | --- |
| ESMC-6B (raw) | 6B | 56.86% | 10.36% | 31.94% |
| **ProtJEPA-600M** | 600M | **58.07%** ns | **11.88%** * | **38.04%** *** |

## Appendix L Limitations: Full Mechanistic Account

1. **Structural suppression.** SCOPe-40 Recall@1: 21.5% vs. 50.4%. Functional distillation competes with structural discriminability; a model optimised for both would require higher structural teacher weights and a separate structural probe.
2. **Human-protein restriction.** All 19,971 proteins from PrimeKG (UniProt/human, HPA, STRING human). Cross-species transfer requires a species-agnostic knowledge graph and tissue atlas.
3. **GO without ancestor propagation.** Slightly inflates Hit@10; information-content-weighted evaluation is future work.
4. **Temporal split.** Pfam-family split does not control for temporal annotation bias; a temporally-stratified split would require annotation timestamp metadata not currently available in PrimeKG.

## Appendix M Failed Ablation Attempts

A 4-teacher subset (ESMC-6B, HGT, BioMedBERT, Disorder) achieved 51.3% GO Hit@10; an 8-teacher variant (excluding HPA and Anatomy) achieved 55.8%, motivating retention of all teachers. A 128-dim predictor head collapsed at batch size 8 (predictor capacity must exceed stable rank *≈*17). VICReg collapsed (Mean Cos *>*0.99) at all tested batch sizes (8–64) and *λ*_cov_ values (1–100).

## Appendix N Full Ablation Few-Shot Tables

Tables N5–N7: (1) No Phase 1 recovers GO few-shot gains but collapses on retrieval (*−*9.72pp vs. ESMC); (2) T1-only degrades monotonically on localisation (45.47%*→*32.05%); (3) Cosine/no-whitening recovers localisation but loses retrieval, confirming whitening enables simultaneous multi-task improvement.

**Table N5.** No Phase 1 ablation (random aggregator) vs. ESMC 600M and ProtJEPA. Bootstrap 95% CI over 10 seeds.

| Label % | GO BP |  |  | Localization |  |  |
| --- | --- | --- | --- | --- | --- | --- |
|  | ESMC | No Ph1 | ProtJEPA | ESMC | No Ph1 | ProtJEPA |
| 1% | 8.85 [7.7,10.0] | 11.18 [10.0,12.2] | **11.88 [10.6,13.0]** | 26.17 [25.1,27.4] | 36.97 [35.9,37.9] | **38.04 [36.6,39.3]** |
| 5% | 11.66 [11.1,12.3] | 13.64 [12.7,14.5] | **14.82 [14.2,15.7]** | 35.01 [34.1,35.7] | 40.77 [40.2,41.3] | **44.62 [44.1,45.1]** |
| 10% | 12.34 [11.7,12.9] | 15.15 [14.6,15.6] | **16.03 [15.6,16.4]** | 38.94 [38.3,39.5] | 42.02 [41.5,42.6] | **46.11 [45.7,46.6]** |
| 50% | 15.19 [14.9,15.4] | 17.51 [17.1,17.9] | **22.85 [22.6,23.1]** | 43.79 [43.6,43.9] | 43.48 [43.2,43.8] | **48.59 [48.4,48.8]** |
| 100% | 15.66 [15.7,15.7] | 18.20 [18.2,18.2] | **25.00 [25.0,25.0]** | 45.47 [45.5,45.5] | 43.91 [43.9,43.9] | **48.51 [48.5,48.5]** |
| GO Hit@10 | 55.27% | 45.55% | **58.07%** | (zero-shot, all $p < 0.0001$) | | |

**Table N6.** T1-only ablation (distilling from ESMC 6B only). Bootstrap 95% CI over 10 seeds.

| Label % | GO BP |  |  | Localization |  |  |
| --- | --- | --- | --- | --- | --- | --- |
|  | ESMC | T1-only | ProtJEPa | ESMC | T1-only | ProtJEPa |
| 1% | 8.85 [7.8,9.9] | 8.16 [6.5,9.8] | **11.88 [10.5,13.0]** | 26.17 [25.1,27.4] | 24.02 [22.9,25.0] | **38.04 [36.5,39.3]** |
| 5% | 11.66 [11.2,12.2] | 10.96 [10.7,11.2] | **14.82 [14.2,15.6]** | 35.01 [34.1,35.7] | 30.11 [29.6,30.7] | **44.62 [44.1,45.1]** |
| 10% | 12.34 [11.7,12.9] | 11.19 [11.0,11.4] | **16.03 [15.6,16.4]** | 38.94 [38.3,39.6] | 30.78 [30.5,31.0] | **46.11 [45.7,46.5]** |
| 50% | 15.19 [14.9,15.5] | 12.72 [12.5,12.9] | **22.85 [22.5,23.2]** | 43.79 [43.6,43.9] | 31.79 [31.5,32.1] | **48.59 [48.4,48.8]** |
| 100% | 15.66 [15.7,15.7] | 13.35 [13.3,13.3] | **25.00 [25.0,25.0]** | 45.47 [45.5,45.5] | 32.05 [32.0,32.0] | **48.51 [48.5,48.5]** |
| GO Hit@10 | 55.27% | 45.62% | **58.07%** | (zero-shot, all $p < 0.0001$) | | |

**Table N7.** Cosine loss without whitening ablation. Bootstrap 95% CI over 10 seeds.

| Label % | GO BP |  |  | Localization |  |  |
| --- | --- | --- | --- | --- | --- | --- |
|  | ESMC | Cosine | ProtJEPa | ESMC | Cosine | ProtJEPa |
| 1% | 8.85 [7.7,10.0] | 11.52 [10.6,12.5] | **11.88 [10.6,13.0]** | 26.17 [25.1,27.2] | 35.90 [34.9,37.0] | **38.04 [36.5,39.3]** |
| 5% | 11.66 [11.1,12.2] | 13.48 [12.8,14.1] | **14.82 [14.2,15.7]** | 35.01 [34.1,35.7] | 39.46 [39.1,39.8] | **44.62 [44.2,45.1]** |
| 10% | 12.34 [11.7,12.9] | 13.87 [13.3,14.3] | **16.03 [15.6,16.4]** | 38.94 [38.2,39.6] | 40.51 [40.2,40.9] | **46.11 [45.7,46.5]** |
| 50% | 15.19 [14.9,15.5] | 15.15 [14.9,15.4] | **22.85 [22.6,23.2]** | 43.79 [43.6,43.9] | 41.31 [41.0,41.6] | **48.59 [48.4,48.8]** |
| 100% | 15.66 [15.7,15.7] | 15.53 [15.5,15.5] | **25.00 [25.0,25.0]** | 45.47 [45.5,45.5] | 41.97 [42.0,42.0] | **48.51 [48.5,48.5]** |
| GO Hit@10 | 55.27% | 44.47% | **58.07%** | (zero-shot, $p < 0.0001$ for all) | | |

